# *Milo:* differential abundance testing on single-cell data using k-NN graphs

**DOI:** 10.1101/2020.11.23.393769

**Authors:** Emma Dann, Neil C. Henderson, Sarah A. Teichmann, Michael D. Morgan, John C. Marioni

## Abstract

Single-cell omic protocols applied to disease, development or mechanistic studies can reveal the emergence of aberrant cell states or changes in differentiation. These perturbations can manifest as a shift in the abundance of cells associated with a biological condition. Current computational workflows for comparative analyses typically use discrete clusters as input when testing for differential abundance between experimental conditions. However, clusters are not always an optimal representation of the biological manifold on which cells lie, especially in the context of continuous differentiation trajectories. To overcome these barriers to discovery, we present *Milo*, a flexible and scalable statistical framework that performs differential abundance testing by assigning cells to partially overlapping neighbourhoods on a k-nearest neighbour graph. Our method samples and refines neighbourhoods across the graph and leverages the flexibility of generalized linear models, making it applicable to a wide range of experimental settings. Using simulations, we show that *Milo* is both robust and sensitive, and can reveal subtle but important cell state perturbations that are obscured by discretizing cells into clusters. We illustrate the power of *Milo* by identifying the perturbed differentiation during ageing of a lineage-biased thymic epithelial precursor state and by uncovering extensive perturbation to multiple lineages in human cirrhotic liver. *Milo* is provided as an open-source R software package with documentation and tutorials at https://github.com/MarioniLab/miloR.

## Introduction

The advent and expansion of high-throughput and high-dimensional single-cell measurements has empowered the discovery of specific cell-state changes associated with disease, development and experimental perturbations. Perturbed cell states can be detected by quantifying shifts in abundance of cell types in response to a biological insult. A common analytical approach for quantitatively identifying such shifts is to ask whether the composition of cells in predefined and discrete clusters differs between experimental conditions [1–5]. However, assigning single-cells to discrete clusters can be problematic, especially in the context of continuous differentiation, developmental or stimulation trajectories, thus limiting the power and resolution of such differential abundance (DA) testing strategies.

Alternative approaches for performing differential abundance testing without requiring clusters to be defined have been proposed for high-throughput mass cytometry data [6]. For example, *Cydar* constructs hyperspheres in the high-dimensional (protein) expression space and asks whether the abundance of cells from different conditions varies in each hypersphere. However, the construction of hyperspheres depends heavily upon the choice of input parameters and upon data pre-processing. More recent developments have proposed strategies for overcoming some of these limitations, but are themselves constrained to pairwise comparisons, as implemented in *DAseq* [7], and thus lack flexibility.

To solve these challenges, we have developed a computational method that performs differential abundance testing without relying on clustering cells into discrete groups. We make use of a common data-structure that is embedded in many single-cell analyses: k-nearest neighbour (k-NN) graphs. We model cellular states as overlapping neighbourhoods on such a graph, which are then used as the basis for differential abundance testing. To account for the non-independence of spatially overlapping neighbourhoods we build upon a previously described strategy to control the spatial False Discovery Rate (FDR) [6].

Our method, which we call *Milo*, leverages the flexibility of generalized linear models (GLM), thus allowing complex experimental designs. Moreover, by modelling cell states as overlapping neighbourhoods, we are able to accurately pinpoint the perturbed cellular states, enabling the identification of the underlying molecular programs. We demonstrate the power of our approach by identifying perturbed cellular states from publicly available datasets in the context of human liver cirrhosis and by uncovering a fate-biased progenitor in the ageing murine thymus. Furthermore, we demonstrate the speed and scalability of our open-source implementation of *Milo*, and demonstrate its superiority to alternative approaches.

## Results

### Modelling cell states as neighbourhoods on a k-NN graph

We propose to model the differences in the abundance of cell states between experimental conditions using graph neighbourhoods (Fig 1). Our computational approach allows overlapping neighbouring regions, which alleviates the principal pitfalls of using discrete clusters for differential abundance testing. We make use of a refined sampling implementation [8], which leads to high coverage of the graph while simultaneously controlling the number of neighbourhoods that need to be tested. For each neighbourhood we then perform hypothesis testing between biological conditions to identify differentially abundant cell states whilst controlling the FDR across the graph neighbourhoods.

**Figure 1:**
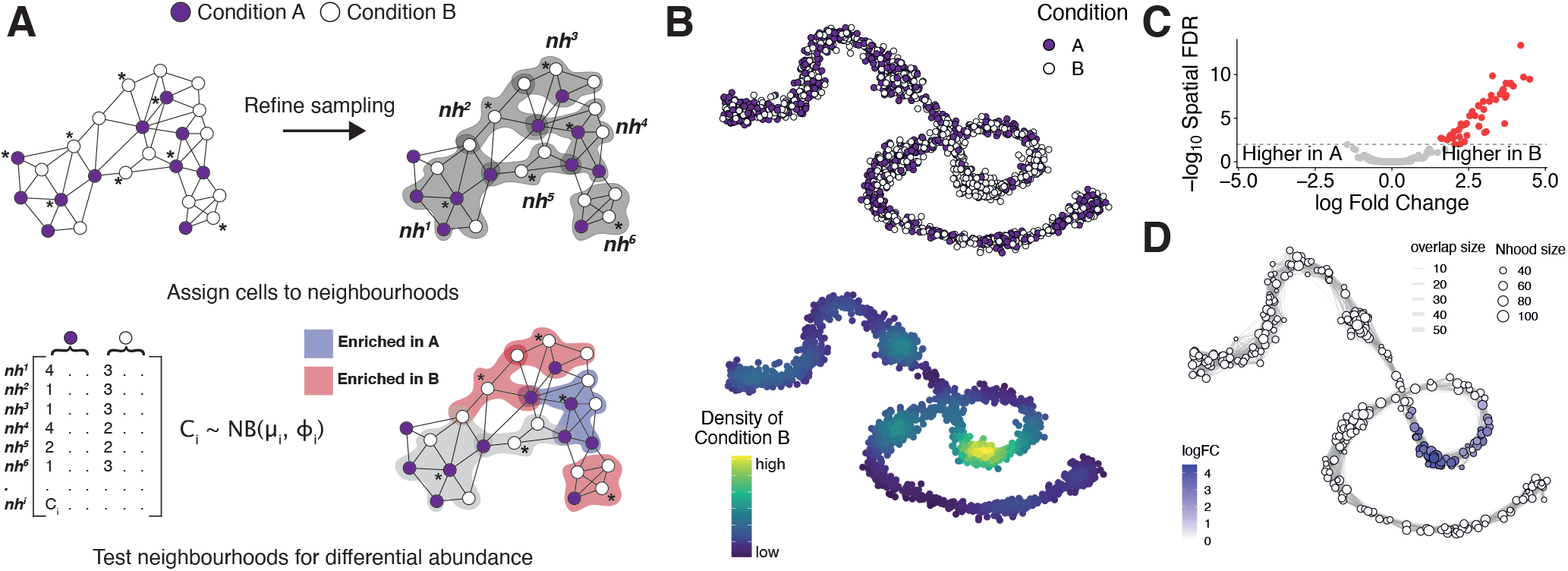
Detecting perturbed cell states as differentially abundant graph neighbourhoods. (A) Schematic of the Milo workflow. Neighbourhoods are defined on index cells, selected using a graph sampling algorithm. Cells are quantified according to the experimental design to generate a counts table. Per-neighbourhood cell counts are modelled using a negative binomial GLM, and hypothesis testing is performed to determine differentially abundant neighbourhoods. (B) A force-directed layout of a k-NN graph representing a simulated continuous trajectory of cells sampled from 2 experimental conditions (top panel - A: purple, B: white, bottom panel - kernel density of cells in condition ‘B’). (C) Hypothesis testing using Milo accurately and specifically detects differentially abundant neighbourhoods (FDR 1%). Red points denote DA neighbourhoods. (D) A graph representation of the results from Milo differential abundance testing. Nodes are neighbourhoods, coloured by their log fold-change. Non-DA neighbourhoods (FDR 1%) are coloured white, and sizes correspond to the number of cells in a neighbourhood. Graph edges depict the number of cells shared between adjacent neighbourhoods. The layout of nodes is determined by the position of the neighbourhood index cell in the force-directed embedding of single cells.

Our method works on a k-NN graph that represents the high-dimensional relationships between single-cells, a common scaffold for many single-cell analyses [1–4] (Fig 1A). The first step in our method is to define a set of representative neighbourhoods on the k-NN graph, where a neighbourhood is defined as the group of cells that are connected to an index cell by an edge in the graph. Consequently, we need to sample a subset of single-cells to use as neighbourhood indices. Adopting a purely random sampling approach means that the number of neighbourhoods required to sample a fixed proportion of cells scales linearly with the total number of index cells (Supp Fig 1B). This leads to an increased multiple testing burden, with the potential to reduce statistical power. To solve this problem we have implemented a refined sampling scheme (Fig 1A) [8]. Concretely, we perform an initial sparse sampling, without replacement, of single-cells and compute the k nearest neighbors for each sampled cell. We then calculate the median position of each set of nearest neighbors and find the nearest cell to this median position. These adjacent cells become the set of indices from which we compute the final set of neighbourhoods. This procedure has three main advantages: (1) fewer, yet more representative, neighbourhoods are selected, as initial random samplings from dense regions of the k-NN graph will often converge to the same index cell (Supp Fig 1A), (2) the representative neighbourhoods include more cells on average (Suppl Fig 1B) and (3) neighbourhood selection is more robust across initializations (Supp Fig 1C).

Next, we count the numbers of cells present in each neighbourhood (per experimental sample) and use these for differential abundance testing between conditions. To incorporate complex experimental designs (e.g., the presence of multiple conditions) we test for differences in abundance using a Negative Binomial GLM framework [9,10]. By doing this, we can borrow information across neighbourhoods, allowing robust estimation of dispersion parameters. We also employ a quasi-likelihood F-statistic [11] for comparing different hypotheses, which has been shown to be powerful in single-cell differential testing [12]. To account for multiple hypothesis testing we use a weighted FDR procedure [13] that accounts for the spatial overlap of neighbourhoods as initially introduced in *Cydar* [6]. We adapt this procedure for a k-NN graph, and weight each hypothesis test P-value by the reciprocal of the kth nearest neighbour distance.

Although the GLM framework allows the incorporation of nuisance covariates, to maximize the power of DA testing, confounding effects should be minimized prior to graph building, for example by applying an appropriate batch integration (practical considerations and demonstrations on how to account for batch effects can be found in the Supplementary Notes and Suppl.Fig.2-3).

To illustrate the *Milo* workflow we generated a simulated trajectory [14] composed of cells sampled from two experimental conditions (‘A’ and ‘B’; Fig1B). Cells in a defined subpopulation of this trajectory were simulated to be more abundant in the ‘B’ condition (Fig 1B); this region of differential abundance is not defined as a distinct cluster by widely-used clustering algorithms (Supp Fig 4). However, applying *Milo* to these simulated data specifically detects that this region contains different abundances of cells from the two conditions (Fig 1C-D).

### Milo out-performs existing methods for differential abundance testing

To illustrate the power and accuracy of *Milo* we first simulated 100 independent continuous trajectories, each consisting of 2000 single-cells, and assigned cells equally to one of 2 conditions: ‘A’ or ‘B’. To simulate a subpopulation of differential abundance we sampled 90% of cells in a specific region of each trajectory from condition ‘B’. Moreover, we assigned cells to one of 3 replicates per condition, thus mimicking a common experimental design. These simulated data sets provide a ground truth against which the performance of differential abundance testing approaches can be compared (Fig 2A).

**Figure 2:**
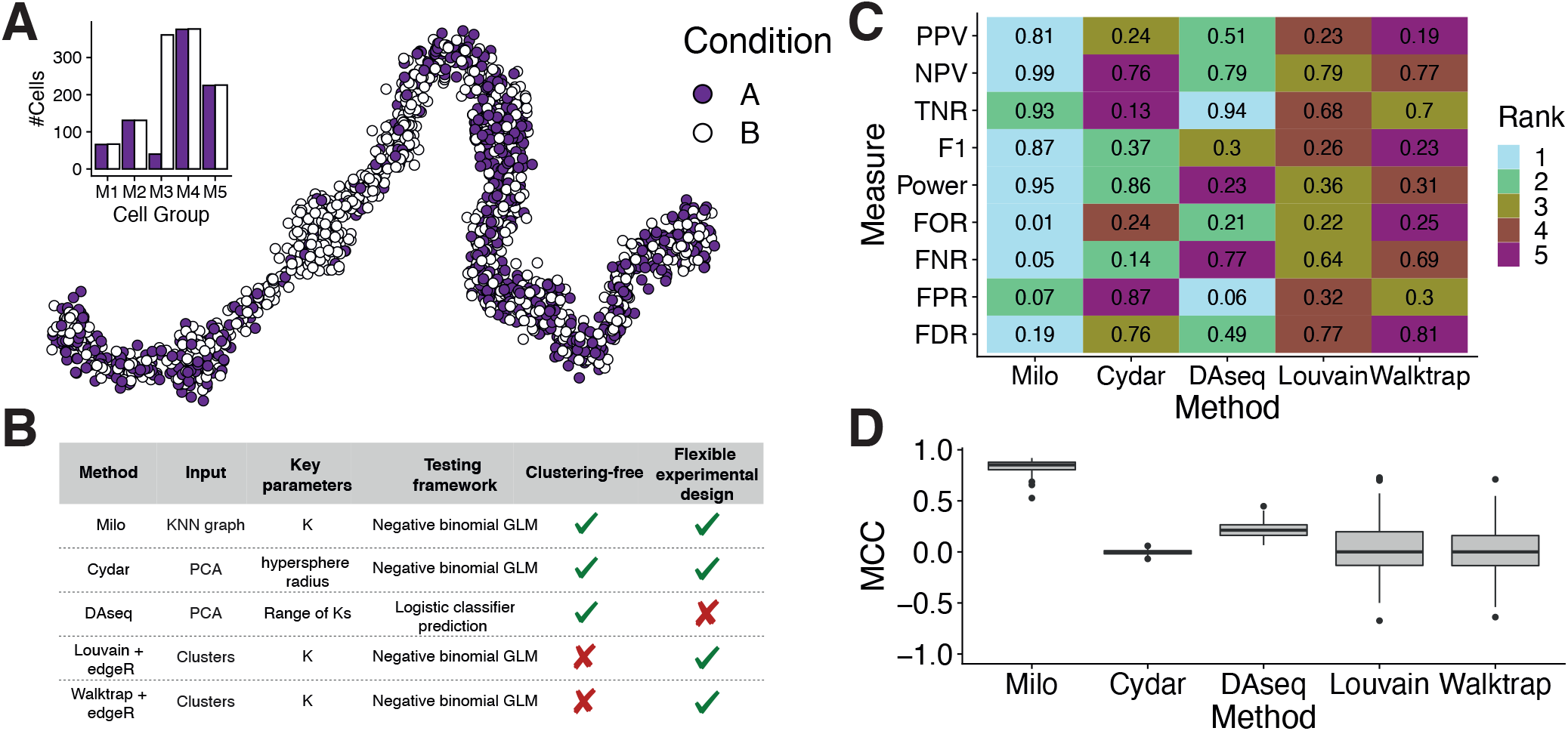
Milo outperforms alternative differential abundance testing approaches. (A) An example simulated trajectory of cells drawn from 5 groups with cells assigned to either conditions ‘A’ (purple points) or ‘B’ (white points). Inset bar plot shows the number of cells (y-axis) assigned to each condition according to the group from which cells were sampled (x-axis). (B) A table describing the different methods compared to Milo, along with the input, key parameters and an overview of the testing framework for each. (C) Rankings of DA testing methods across a number of measures to determine performance. Each box is coloured by the ranking of each measure for each method, where a rank of 1 indicates the best performance and 5 indicates the worst across 100 simulated data sets. Ranks are calculated from the mean value across 100 simulated data sets; mean values are shown. PPV: positive predictive value, NPV: negative predictive value, TNR: true negative rate, F1: F1 score, FOR: false omission rate, FNR: false negative rate, FPR: false positive rate, FDR: false discovery rate. (D) The Matthews correlation coefficient assesses the performance of each method by integrating across multiple performance measures. Box plots show the MCC across 100 independent simulations for each method.

As well as *Milo*, we applied two methods designed for differential abundance testing using single-cell data to these simulated datasets: *Cydar*, originally designed to model differential abundance in mass cytometry data [6], and *DAseq*, which utilises a logistic classifier to predict which cells are from single-cell DA subpopulations represented by a reduced dimensional space [7] (Fig 2B). In addition, we applied the current best-practise analysis strategy for single-cell analysis: graph-clustering followed by differential abundance testing of clusters between conditions. For this approach we applied 2 commonly used community detection algorithms: Louvain [15] and Walktrap [16]. We modelled the differential abundance of clusters from these algorithms using a NB GLM, as implemented in *edgeR* [9]. To ensure comparability between methods we used the same reduced dimensional space as the input for all methods and the same parameter values, where these were shared, e.g. the value of ‘k’ for k-NN graph building. Where parameters were specific to a method, we made use of the recommended practise by the method developers to select an appropriate value (Supp Table 1).

For each simulated dataset we computed the confusion matrix of each method against the ground truth, and calculated a number of common summary statistics (Fig 2C, Supp Fig 5), enabling an assessment of how well each method performs across a variety of metrics. To generate a single value for comparing methods, and integrate across the four categories of a confusion matrix, we also calculated the Matthews correlation coefficient (MCC) [17] (Fig 2D). The MCC takes values between 1 (highly consistent) and -1 (highly inconsistent), thus providing an intuitive assessment of method performance. Across all 100 simulations we found that *Milo* out-performed all other methods, including both clustering methods, demonstrating the additional gains of modelling cell states using overlapping neighbourhoods (Fig 2C, Supp Fig 5). This was confirmed when examining the MCC, where we observed that *Milo* yielded the highest median correlation (0.85) and lowest variance (Fig 2D). Conversely, the clustering-based methods resulted in highly variable MCC values, illustrating the sensitivity of these approaches to the input data set. In sum, our simulations demonstrate that *Milo* circumvents a common bottleneck in single-cell analyses: the need to perform iterative rounds of community detection to achieve an optimal clustering prior to differential abundance testing.

### Milo is fast and scalable

The benchmarking dataset is fairly typical in size for current single-cell experiments. However, moving forward, the number of cells assayed is likely to increase with advances in experimental sample multiplexing [18,19]. As such, we tested the scalability of the *Milo* workflow, and profiled the memory usage across multiple steps. For this we ran *Milo* on 3 published datasets of differing sizes from ∼2000 to ∼130,000 cells, representing differences in both biological and experimental complexity [2–4], as well as a dataset of 200,000 simulated single-cells from a linear trajectory (see Methods). Using these 4 data sets we measured the amount of time required to execute the *Milo* workflow from graph-building through to differential abundance testing (Fig 3A). In parallel, we profiled the amount of memory used across the entire workflow (Fig 3B) and at each defined step (Supp Fig 6). Notably, the amount of time taken increased linearly with the total size of the data set (Fig 3A), which for a large set of 200k cells was less than 90 minutes. Moreover, the total memory usage across all steps of the *Milo* workflow scaled primarily with the size of the input dataset (Fig 3B), indicating that the complexity and composition of the single-cells largely determines the memory requirements (Supp Fig 6). Importantly, these memory requirements are within the resources of common desktop computers (i.e. <16GB). This benchmarking analysis demonstrates that *Milo* is able to perform differential abundance analysis in large and complex datasets at a scale and speed that is feasible on a desktop computer.

**Figure 3:**
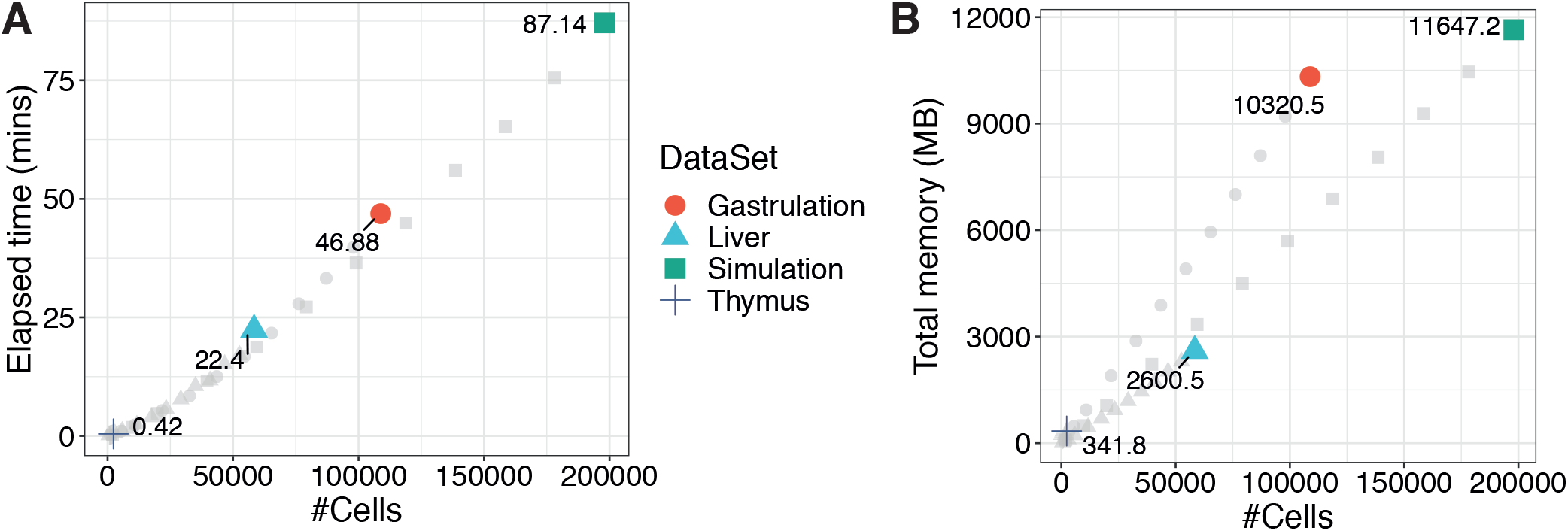
Milo efficiently scales to large data sets. (A) Run time (y-axis) of the Milo workflow from graph building to differential abundance testing. Each point represents a down-sampled dataset, denoted by shape. Coloured points show the total number of cells in the full dataset labelled by the elapsed system time (mins). (B) Total memory usage (y-axis) across the Milo workflow. Each point represents a down-sampled dataset, denoted by shape. Coloured points are the full datasets labelled with the total memory usage (megabytes).

### Milo identifies the decline of a fate-biased epithelial precursor in the ageing mouse thymus

To demonstrate the utility of *Milo* in a real-world setting we applied it to a single-cell RNA-seq dataset of mouse thymic epithelial cells (TEC) sampled across the first year of mouse life, which were previously clustered into 9 distinct TEC subtypes (Fig 4A) [3]. These data, generated using plate-based SMART-seq2, consist of 2327 single-cells equally sampled from mice at 5 different ages: 1, 4, 16, 32 and 52 weeks old (Fig 4B). Moreover, the experimental design included 5 replicate experimental samples of cells for each age. The goal of the study was to identify TEC subtypes that change in frequency during natural ageing.

**Figure 4:**
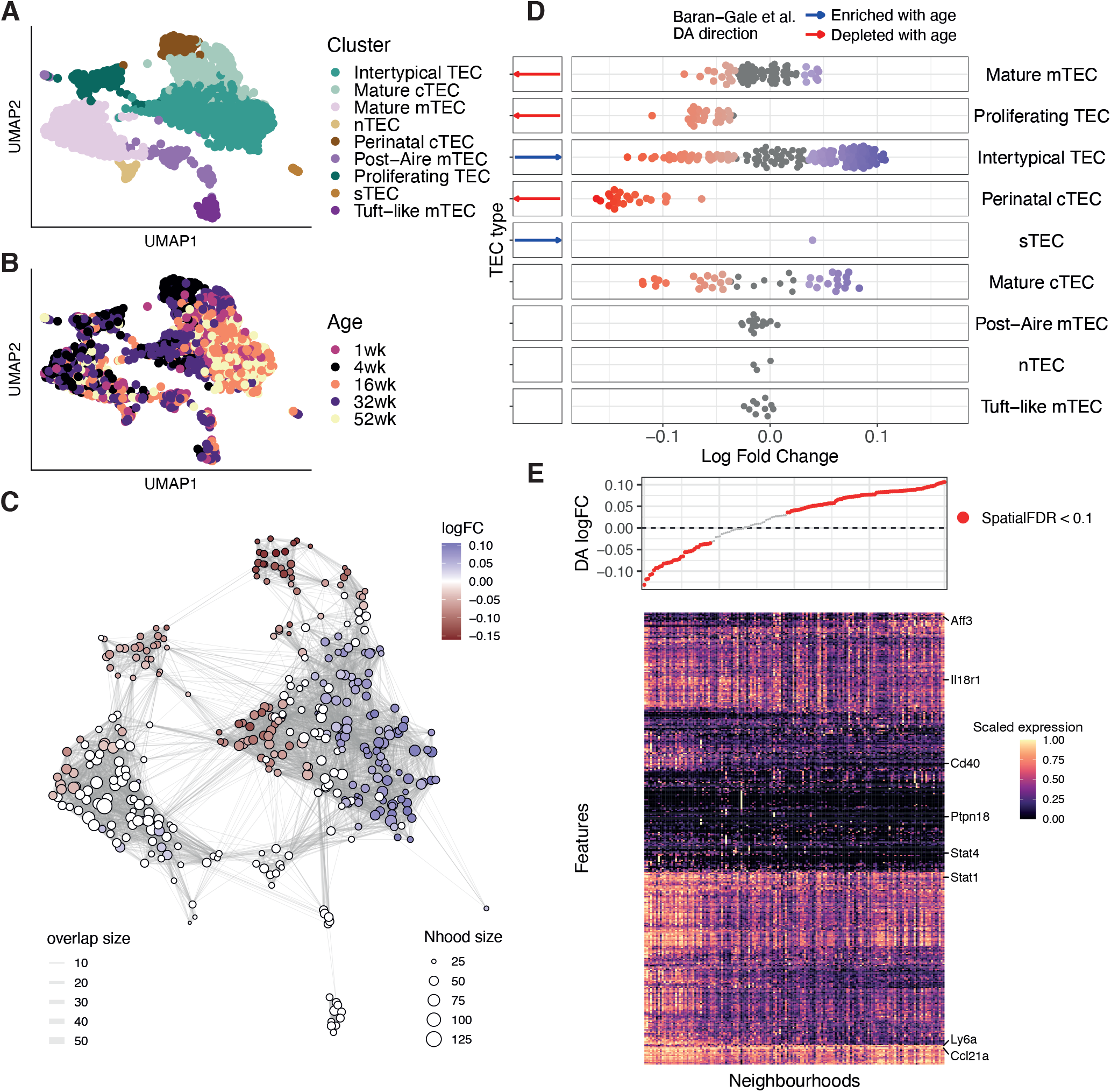
Milo identifies the decline of a fate-biased precursor in the ageing mouse thymus. (A-B) A UMAP of single thymic epithelial cells sampled from mice aged 1-52 weeks old. Points are labelled according to their annotation in Baran-Gale et al. 2020 (A) and mouse age (B) (C) A graph representation of the results from Milo differential abundance testing. Nodes are neighbourhoods, coloured by their log fold change across ages. Non-DA neighbourhoods (FDR 10%) are coloured white, and sizes correspond to the number of cells in a neighbourhood. Graph edges depict the number of cells shared between adjacent neighbourhoods. The layout of nodes is determined by the position of the neighbourhood index cell in the UMAP embedding of single cells. (D) Beeswarm plot showing the distribution of log-fold change across age in neighbourhoods containing cells from different cell type clusters. DA neighbourhoods at FDR 10% are coloured. Cell types detected as DA through clustering by Baran-Gale et al. (2020) are annotated in the left side bar. (E) A heatmap of genes differentially expressed between DA neighbourhoods in the Intertypical TEC cluster. Each column is a neighbourhood and rows are differentially expressed genes (FDR 1%). Expression values for each gene are scaled between 0 and 1. The top panel denotes the neighbourhood DA log fold-change.

To this end, we first constructed a k-NN graph, before assigning cells to 363 neighbourhoods, which were then used to test for differential abundance of TEC states across time. At a 10% FDR, we identified 217 DA neighbourhoods (112 showed a decreased abundance with age, 105 an increased abundance with age) spanning multiple TEC states (Fig 4C). We compared our results to those generated in the original publication, which demonstrated that we were able to identify the same DA states (Fig 4D), including changes in the abundance of the ‘sTEC’ population, which consisted of just 24 cells. Moreover, whilst we recovered the previously reported accumulation of Intertypical TEC with age, we also identified a novel subset of these cells that were depleted with age (Fig 4C-D).

We have previously shown that Intertypical TEC represent an adult progenitor of medullary TEC (mTEC) [3]. To understand the function of the novel sub-state of Intertypical TEC identified using *Milo* we performed marker gene expression identification between the Intertypical TEC in neighbourhoods enriched or depleted in younger mice (FDR 1%; Fig4E). This analysis indicated that the cells from younger mice up-regulated multiple cytokine response genes (e.g. *Stat1, Stat4, Aff3*) (Fig 4E), illustrated by the enriched Gene Ontology term GO:0034097 ‘response to cytokine’ (enrichment adjusted p-value=2.48×10^−3^). Cytokine signalling is key to mTEC differentiation [20,21], indicating that these TEC from younger mice might be differentiating more efficiently to the mTEC lineage. The discovery that medullary-biased Intertypical TEC are less abundant with age was corroborated by our original study utilising a much larger data set of ∼90,000 single-cells coupled with lineage-tracing [3]. Therefore, these analyses demonstrate the sensitivity of *Milo* by identifying that a mTEC progenitor state is depleted with age, a finding that was not resolved using clustering approaches.

### Milo identifies compositional disorder in cirrhotic human liver

To demonstrate the applicability of our method in multiple biological contexts, we next applied *Milo* to a large dataset of hepatic cells isolated from 5 healthy and 5 cirrhotic human livers [2]. The original study assigned cells to multiple lineages, including immune, endothelial and mesenchymal cells (Fig 5A-B). A key goal of the study was to ask whether different cell types were differentially abundant between experimental samples taken from healthy and cirrhotic tissue. In the original study, cells from each lineage were sub-clustered and these sub-clusters were interrogated using a Poisson GLM to identify whether there were differential contributions from cirrhotic and healthy donors.

**Figure 5:**
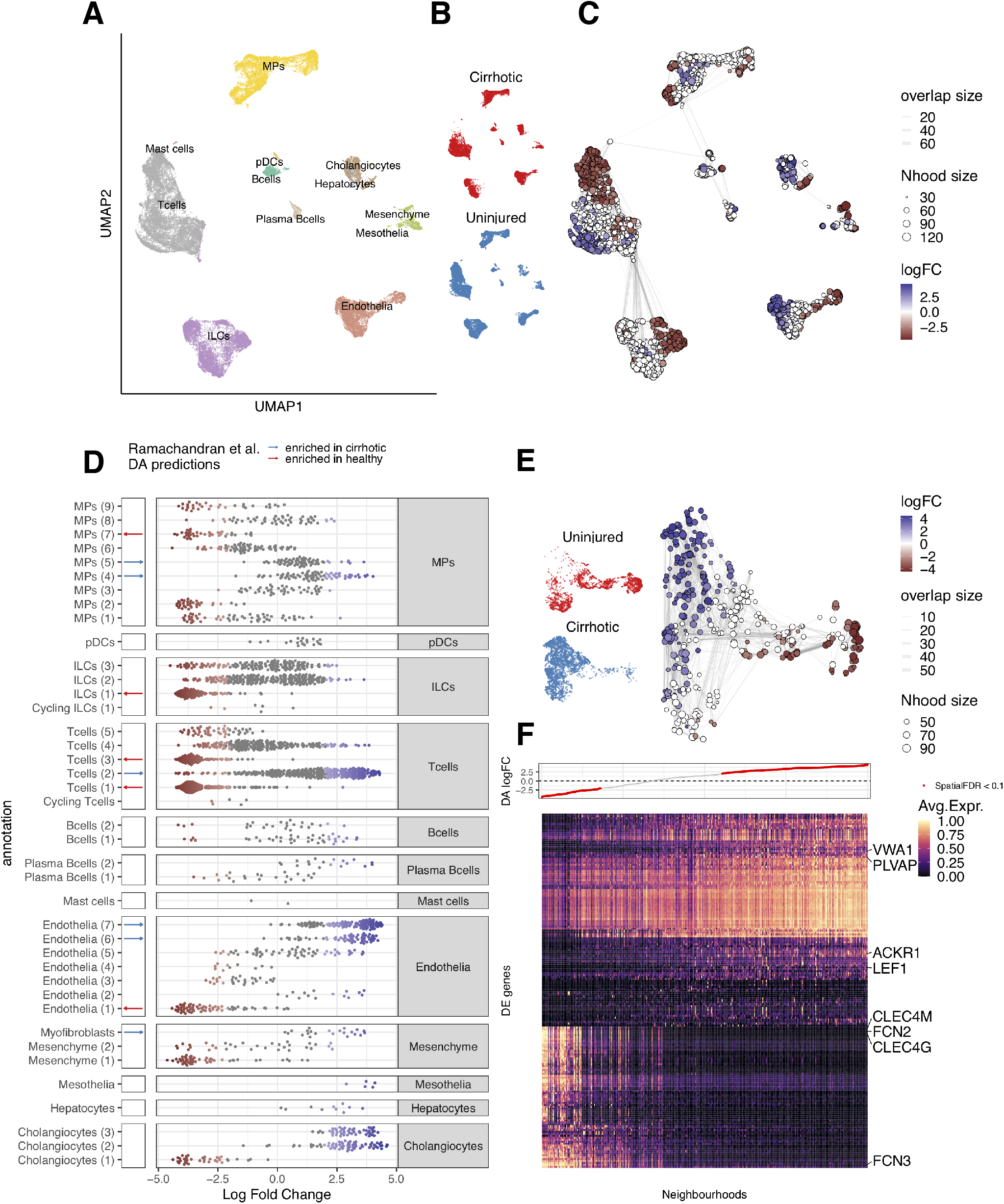
Milo identifies the compositional disorder in cirrhotic liver. (A-B) UMAP embedding of 58358 cells from healthy (n = 5) and cirrhotic (n = 5) human livers. Cells are colored by cellular lineage (A) and injury condition (B) (C) Graph representation of neighbourhoods identified by Milo. Nodes are neighbourhoods, coloured by their log fold change between cirrhotic and healthy samples. Non-DA neighbourhoods (FDR 10%) are coloured white, and sizes correspond to the number of cells in a neighbourhood. Graph edges depict the number of cells shared between adjacent neighbourhoods. The layout of nodes is determined by the position of the neighbourhood index cell in the UMAP embedding of single cells. (D) Beeswarm plot showing the distribution of log-fold change in abundance between conditions in neighbourhoods from different cell type clusters. DA neighbourhoods at FDR 10% are coloured. Cell types detected as DA through clustering by Ramachandran et al. (2019) are annotated in the left side bar. (E) UMAP embedding and graph representation of neighbourhoods of 7995 cells from endothelial lineage. (F) Heatmap showing average neighbourhood expression of genes differentially expressed between DA neighbourhoods in the endothelial lineage (572 genes). Expression values for each gene are scaled between 0 and 1. The top panel denotes the neighbourhood DA log fold-change.

To explore whether more subtle differences could be detected, we applied *Milo* analysis to 2696 neighbourhoods spanning the k-NN graph and identified 1404 neighbourhoods with differential abundance (10% FDR; Fig 5C). To assess performance, we compared DA results with those from the compositional analysis performed by Ramachandran et al. [2]. *Milo* recovered DA neighbourhoods in all clusters identified as differentially abundant between cirrhotic or uninjured tissue in the original study (Fig 5D).

Moreover, *Milo* identified multiple groups of neighbourhoods within the same pre-defined sub-clusters that showed opposing directions of differential abundance between the control and cirrhotic liver experimental samples (Fig 5D). In other words, within a sub-cluster, some neighbourhoods were enriched for control experimental samples whilst others were enriched for disease experimental samples. These patterns, exemplified by the T cell (2) and the endothelial (5) compartments were obscured in the previous study due to the reliance on pre-clustering (Fig 5D).

To further explore the biological meaning of these neighbourhoods, we first focused on the hepatic endothelial cells, where we resolved disease specific subpopulations at higher resolution than was possible by clustering-based analysis (Fig. 5D). *Milo* identified a gradient of changes in neighbourhood abundance across this compartment, suggestive of a continuous transition between healthy and diseased cell states (Fig 5E). To identify gene expression signatures associated with this change, we performed differential expression analysis between cells in DA neighbourhoods with positive and negative log fold changes, identifying 83 differentially expressed genes (FDR 10%; Methods) (Fig5F). In the cirrhosis-enriched neighbourhoods, we recovered over-expression of known markers of scar-associated endothelium, including *ACKR1, PLVAP* and *VWA1* (Fig. 5F) [2]. We also recovered over-expression of genes associated with regulation of leukocyte recruitment, confirming the validated immunomodulatory phenotype displayed by scar-associated tissue (Supp Fig 7A) [22]. In addition, cirrhotic endothelium displays down-regulation of genes involved in response to infection, endocytosis and immune complex clearance, including *FCN2, FCN3*, and *FCGR2B* (Supp Fig 7B), which have been suggested as an additional component of cirrhosis-associated immune dysfunction [23,24].

*Milo* also identified strong DA between healthy and cirrhotic cells in lineages that were unexplored in the original study, such as the cholangiocyte compartment (Fig 5D). Cholangiocytes are epithelial cells that line a three-dimensional network of bile ducts known as the biliary tree, and cholangiocyte proliferation can be induced by a broad range of liver injuries, in a process termed the ductular reaction [25]. However, the gene signatures associated with this process in human cirrhosis are largely unexplored. Indeed, *Milo* recovered an enrichment of disease-specific cholangiocytes (Supp Fig 7C-D), and differential gene expression analysis detected strong over-expression of genes associated with calcium signalling (Supp Fig 7E-F), a signalling pathway frequently dysregulated in liver disease and a potential target for clinical intervention [26,27].

These analyses demonstrate the potential of using DA subpopulations detected by *Milo* to recover known and novel signatures of disease-specific cell states.

## Discussion

Given the increasing number of complex single-cell datasets where multiple conditions are assayed [18,19], *Milo* tackles a key problem: robustly determining sets of cells that are differentially abundant between conditions without relying on pre-existing sets of clusters. Moreover, *Milo* is fully interoperable with established single-cell analysis workflows and is implemented as an open-source R software package [28] with documentation and tutorials at https://github.com/MarioniLab/miloR.

The definition of neighbourhoods, as implemented in *Milo*, overcomes the main limitations of standard-of-practice clustering-based DA analysis, whilst utilising a common data-structure in single-cell analysis - graphs. A strength of our approach is that it is applicable to a wide range of datasets with vastly different topologies, including gradual state transitions, thus removing the need for time-consuming iterative sub-clustering and identifying subtle differences in differential abundance that would otherwise be obscured (Fig 5D).

Recently, other clustering-free methods have been proposed to detect compositional differences between experimental conditions [7,29]. However, these are most suitable for pairwise comparisons between two biological conditions, and cannot be easily extended to detect changes across continuous conditions (age, time points) or multifactorial conditions. By modelling cell counts with a NB GLM, *Milo* can incorporate arbitrarily complex experimental designs as demonstrated by our application of Milo to detect compositional changes in the ageing mouse thymus (Fig 4) and across early embryonic development in mice (Supp Fig 3). Moreover, we show how nuisance technical covariates can be included in the GLM model to increase the power of DA testing in the presence of batch effects (Supp Fig 2-3).

Although we have addressed several important challenges, *Milo* is not free of limitations. Firstly, the testing framework requires a replicated experimental design to estimate the dispersions of counts for each condition. Whilst this is not strictly a limitation of *Milo*, it reflects the importance of properly replicated experimental design in single-cell experiments. A potential solution would be to use a mixed effects model utilising random intercepts. Secondly, the detection of DA subpopulations by Milo requires a k-NN graph that reflects the true cell-cell similarities in the phenotypic manifold; a limitation shared with all DA methods that work on graphs or reduced dimensional spaces [30]. Additionally, while *Milo* can account for artefacts such as batch effects during DA testing, we show that optimal results are achieved when batch correction is performed prior to graph construction (Suppl. Note 2, Suppl. Fig. 1, Suppl. Fig. 2). Thirdly, cells in a single neighbourhood do not necessarily represent a unique biological subpopulation; a cellular state might span multiple neighbourhoods. Accordingly, we search for marker genes of DA states by aggregating cells in adjacent and concordantly DA neighbourhoods (Fig. 4E, 5F). One challenge of this approach is that rare cell states may be represented by a small subset of neighbourhoods, thus making identification of marker genes challenging. To overcome this problem one can either choose a smaller value of *k* or alternatively construct a graph on cells from a particular lineage of interest.

Following the generation of reference single-cell atlases for multiple organisms and tissues, an increasing number of studies now focus on quantifying how cell populations are perturbed in disease, ageing, and development, using, for example, large scaled pooled CRISPR screens [31–33]. We envision that *Milo* will see use in all of these contexts. By leveraging a cell-cell similarity structure, *Milo* is also applicable to single-cell assays other than scRNA-seq, including multi-omic assays [34–38]. Thus, *Milo* has the potential to facilitate the discovery of fundamental biological and medically important processes across multiple layers of molecular regulation when they are assayed at single-cell resolution.

## Methods

### Milo

Milo detects sets of cells that are differentially abundant between conditions by modelling counts of cells in neighbourhoods on a k-NN graph. The workflow includes the following steps:

#### (A) Construction of the k-NN graph

Similar to many other tasks in single-cell analysis, *Milo* uses a k-NN graph computed based on similarities in gene expression space as a representation of the phenotypic manifold in which cells lie. We assume that the k-NN graph is a faithful representation of the single cell phenotypes. Therefore, any batch effect should be corrected prior to graph building to maximize the power of DA testing. In addition, nuisance covariates can be incorporated in the experimental design of the NB GLM framework (practical considerations and demonstrations on how to account for batch effects can be found in the Supplementary Notes and Supp Fig 2-3). Throughout this paper, we build the k-NN graph based on similarity in reduced principal component (PC) space.

#### (B) Definition of cell neighbourhoods

We define the neighbourhood *n*_*j*_ of cell *j* as the group of cells that are connected to *j* by an edge in the graph. We refer to *j* as the index of the neighbourhood. In order to define a representative subset of neighbourhoods that span the whole k-NN graph, we implement a previously adopted algorithm to sample the index cells in a graph [8,39] (See Supplementary Note 1.1.2 for a detailed description).

#### (C) Counting cells in neighbourhoods

For each neighbourhood we count the number of cells from each experimental sample, constructing a neighbourhood x experimental sample count matrix.

#### (D) Testing for differential abundance in neighbourhoods

To test for differential abundance, we analyse neighbourhood counts using the quasi-likelihood (QL) method in `edgeR`, similarly to the implementation in *Cydar* [6]. We fit a NB GLM to the counts for each neighbourhood and use the QL F-test with a specified contrast to compute a P value for each neighbourhood. Details of the statistical framework are provided in Supplementary Note 1.1.3

#### (E) Controlling the Spatial FDR in neighbourhoods

To control for multiple testing, we adapt the Spatial FDR method introduced by *Cydar* [6]. The Spatial FDR can be interpreted as the proportion of the union of neighbourhoods that is occupied by false-positive neighbourhoods. To control the spatial FDR in the k-NN graph, we apply a weighted version of the Benjamini-Hochberg (BH) method, where P values are weighted by the reciprocal of the neighbourhood connectivity. As a measure of neighbourhood connectivity, we use the Euclidean distance to the k-th nearest neighbour of the index cell for each neighbourhood.

A full description of *Milo* can be found in Supplementary Notes.

### Visualization of DA neighbourhoods

To visualize results from differential analysis on neighbourhoods, we construct an abstracted graph, where nodes represent neighbourhoods and edges represent the number of cells in common between neighbourhoods. The size of nodes represents the number of cells in the neighbourhood. The position of nodes is determined by the position of the sampled index cell in the single-cell UMAP, to allow qualitative comparison with the single cell embedding.

### Mouse thymus analysis

Single-cell data are available from ArrayExpress (accession E-MTAB-8560), additional meta-data were acquired from Baran-Gale *et al*. [3] including cluster identity and highly variable genes (HVGs). The dataset consists of 2327 single thymic epithelial cells that passed QC (see [3] for details). Log-normalized gene expression values were used as input, along with 4906 HVGs, to estimate the first 50 principal components using a randomized PCA implemented in the R package `irlba`, the first 40 of which were used for k-NN graph building (k=21) and UMAP embedding. The refined sampling, using an initial random sampling of 30% of cells, identified 363 neighbourhoods. Differential abundance testing used the mouse age as a linear predictor variable, thus log fold changes are interpreted as the per-week linear change in neighbourhood abundance. Neighbourhood cluster identity was assigned by taking the most abundant cluster identity amongst neighbourhood cells.

Differential expression (DE) testing was performed on cells within neighbourhoods containing a majority of cells from the Intertypical TEC cluster. This subset of neighbourhoods was aggregated into 2 groups based on similarity of log fold change direction and number of overlapping cells (≥10 cells). DE testing was performed comparing the log normalized gene expression of neighbourhood cells between the more and less abundant groups using a linear model implemented in the Bioconductor [40,41] package `limma` [42], using 1% FDR. Gene Ontology Biological Process term analysis was performed on the 448 DE genes (adj. P-value < 0.01) using the R package `enrichR` [43].

### Liver cirrhosis analysis

The dataset including cell type annotations was downloaded from https://datashare.is.ed.ac.uk/handle/10283/3433 (GEO accession: GSE136103 [2]). The dataset comprises 58358 cells, obtained from 5 healthy and 5 cirrhotic liver samples. Following the pre-processing steps from the original publication, dimensionality reduction with PCA was performed on the 3000 top highly variable genes (HVGs), calculated using `modelGeneVar` and `getTopHVGs` from the R package `scran` [44], and the top 11 PCs were retained for k-NN graph building and UMAP embedding. Refined sampling identified 2676 neighbourhoods (k=30, initial proportion of sampled cells = 0.05). We run *Milo* to test for DA between cirrhotic and healthy experimental samples. To assign cell type annotations to neighbourhoods, we take the most frequent annotation between cells in each neighbourhood. Neighbourhoods are generally homogeneous, with an average of 80% of cells belonging to the most abundant cell type label.

For the focused analysis on the endothelial and cholangiocyte lineages, DE testing was performed on the subset of neighbourhoods from the selected lineage. Neighbourhoods were aggregated into 2 groups based on similarity of log fold change direction. DE testing was performed summing the gene expression counts for each experimental sample and neighbourhood group between the more and less abundant groups using the quasi-likelihood test implemented in `edgeR` [9], using 10% FDR. GO term analysis was performed on the significant DE genes using the R package `clusterProfiler` [45].

### Mouse gastrulation data

The raw count matrix and batch corrected PCA matrix were downloaded via the R package MouseGastrulationData [46]. To construct the uncorrected k-NN graph, raw counts were log transformed and PCs were computed on the 5000 top variable genes. Refined sampling identified 11895 neighbourhoods in the uncorrected graph and 8451 neighbourhoods in the MNN corrected graph (k = 30, initial proportion of sampled cells = 0.1).

### Scalability analysis

We assessed the scalability of *Milo* by profiling the time taken to execute the workflow, starting with the k-NN graph building step and concluding with the differential abundance testing. We simulated a dataset of 200000 single-cells using the `dyntoy` package implemented in R [14]. With this large simulation we down-sampled to specific proportions, ranging from 1 to 100%, and recorded the elapsed system time to complete the *Milo* workflow using the `system.time` function in R [28]. In addition, we performed an equivalent analysis using the published datasets included in this manuscript: mouse thymus [3], human liver [2], and mouse gastrulation [4]. All timings are reported in minutes.

To assess the memory usage of the *Milo* workflow we made use of the `Rprof` function in R to record the total amount of memory used at each step. We followed the same approach as above, down-sampling simulated and published datasets from 1 to 100% of the total cell numbers. All memory usage is reported in megabytes (MB).

For both the system timing and memory usage we ran the simulated and published datasets down-sampling analyses on a single node of the high-performance computing (HPC) cluster at the Cancer Research UK - Cambridge Institute. Each node has 2x Intel Xeon E5-2698 2.20Ghz processors with 40 cores per node and 384GB DDR4 memory; cluster jobs were run using a single core.

### Batch effect analysis

#### Simulated data

we simulated a dataset representing a continuous trajectory of 5000 cells using the R package `dyntoy` [14]. We assigned cells to one of 6 experimental samples and samples to one of 2 conditions. In a specific region of the trajectory we assigned 80% of cells to condition ‘B’ and 20% to condition ‘A’, simulating differential abundance between conditions. A batch effect was incorporated by adding a gaussian random vector to the expression profiles of all cells in two of six samples. We performed batch correction using the MNN method, as implemented in the R package `batchelor` by the function `fastMNN`, using default parameters [47]. Refined sampling identified 298 neighbourhoods in the uncorrected graph and 317 neighbourhoods in the MNN corrected graph (k = 10, initial proportion of sampled cells = 0.1). We ran *Milo* to test for DA between conditions, with and without accounting for the batch effect in the experimental design (`design = ∼ condition` or `design = ∼ batch + condition`).

### Mouse gastrulation data

We ran *Milo* to test for DA with 3 alternative experimental designs: (A) test for DA across developmental time points (`design = ∼ time point`), (B) test for DA across developmental time points, accounting for the sequencing batch (`design = ∼ seq. batch + time point`) and (C) test for DA between sequencing batches (`design = ∼ seq. batch`).

## Code and data availability

*Milo* is implemented as an open-source package in R: https://github.com/MarioniLab/miloR. Code used to generate figures and perform analyses can be found at https://github.com/MarioniLab/milo_analysis_2020.

## Acknowledgments & Funding

We thank Shila Ghazanfar for feedback on the method, Natsuhiko Kumasaka for comments on the manuscript, Chenqu Suo, Veronika Kedlian and Rasa Elmentaite for feedback on the software package. JCM acknowledges core funding from EMBL and core funding from Cancer Research UK (C9545/A29580), which supports MDM. ED and SAT acknowledge Wellcome Sanger core funding (WT206194). NCH is supported by a Wellcome Trust Senior Research Fellowship in Clinical Science (ref. 219542/Z/19/Z), Medical Research Council, and a Chan Zuckerberg Initiative Seed Network Grant.

## Author contributions

ED, MDM & JCM conceived the method idea. ED & MDM developed the method, wrote the code and performed analyses. ED, MDM, SAT & NCH interpreted the results. ED, MDM, SAT, NCH & JCM wrote and approved the manuscript. MDM & JCM oversaw the project.

