## Supplemental files for "*Milo:* differential abundance testing on single-cell data using k-NN graphs"

### Supplementary information for **Differential cell-state abundance testing using k-NN graphs with *Milo***

**References** **5**

### 1 Supplementary notes

#### 1.1 Description of *Milo*

##### 1.1.1 Building the k-NN graph

Similarly to many other tasks in single-cell analysis, *Milo* uses a k-NN graph computed based on similarities in gene expression space as a representation of the phenotypic manifold in which cells lie. While *Milo* can be used on graphs built with different similarity kernels, here we compute the graph as follows: a gene expression matrix of $M$ cells is projected onto the first $d$ principal components (PCs) to obtain a $M \times d$ matrix $X_{PC}$. Then, for each cell $j$, the Euclidean distances to its $k$ nearest neighbors in $X_{PC}$ are computed and stored in a $M \times M$ adjacency matrix. Then, $D$ is made symmetrical, such that cells $j$ and $l$ are nearest neighbors (i.e. connected by an edge) if either $j$ is a nearest neighbor of $l$ or $l$ is a nearest neighbor of $j$. The k-NN graph is encoded by the undirected symmetric version of $\tilde{D}$ of $D$, where each cell has at least K nearest neighbors.

##### 1.1.2 Definition of cell neighbourhoods and index sampling algorithm

We define the neighbourhood $n_i$ of cell $j$ as the group of cells that are connected to $j$ by an edge in the graph. Here, $i = 1, 2, ..., N$ indexes the sampled neighbourhoods, while $j = 1, 2, ..., M$ indexes the single-cells in the graph, so that $N \leq M$. Formally, a cell $l$ belongs to neighbourhood $n_i$ if $\tilde{D}_{j,l} > 0$. We refer to $j$ as the index cell of the neighbourhood.

In order to define a representative subset of neighbourhoods that span the whole k-NN graph, we implement a previously adopted algorithm to sample the index cells in a graph [1,2]. Briefly, we start by randomly sampling $p \cdot M$ cells from the dataset, where $p \in [0, 1]$ (we use $p = 0.1$ by default). Given the reduced dimension matrix used for graph construction $X_{PC}$, for each sampled cell we consider its $k$ nearest neighbors $l = 1, 2, ..., k$ with PC profiles $x_1, x_2, ..., x_k$. We measure the mean PC profile $\bar{x}$ for the $k$ cells and search for the cell $j$ such that the Euclidean distance between $x_j$ and $\bar{x}$ is minimized. This yields a set of $N \leq p \cdot M$ index cells that are used to define neighbourhoods.

##### 1.1.3 Testing for differential abundance in neighbourhoods

*Milo* builds upon the framework for differential abundance testing implemented by *Cydar* [3]. In this section, we briefly describe the statistical model and adaptations to the k-NN graph setting.

**Quasi-likelihood negative binomial generalized linear models** We consider a neighbourhood $n$ with cell counts $y_{ns}$ for each experimental sample $s$. The counts are modelled by the negative binomial (NB) distribution, as it is supported over all non-negative integers and can accurately model both small and large cell counts. For such non-Normally distributed data we use generalized-linear models (GLMs) as an extension of classic linear models that can accomodate complex experimental designs. We therefore assume that

$$y_{ns} \sim NB(\mu_{ns}, \phi_n),$$

where $\mu_{ns}$ is the mean and $\phi_n$ is the NB dispersion parameter. The expected count value for neighbourhood $n$ in experimental sample $s$, $\mu_{ns}$ is given by

$$\mu_{ns} = \lambda_{ns} M_s$$

where $\lambda_{ns}$ is the proportion of cells belonging to experimental sample $s$ in $n$ and $M_s$ is the total number of cells of $s$. In practice, $\lambda_{ns}$ represents the biological variability that can be affected by treatment condition, age or any biological covariate of interest. We use a log-linear model to represent the influence of the biological condition on the expected counts in neighbourhoods:

$$log\ \mu_{ns} = \sum_{g=1}^{G} x_{sg} \beta_{ng} + log\ M_s$$

where $x_{sg}$ is the covariate vector indicating the condition applied to sample $s$ and $\beta_{ng}$ is the regression coefficient by which the covariate effects are mediated for neighbourhood $n$.

Estimation of $\beta_{ng}$ for each $n$ and $g$ is performed by fitting the GLM to the count data for each neighbourhood, i.e. by estimating the dispersion $\phi_n$ that models the variability of cell counts for replicate samples for each neighbourhood. Dispersion estimation is performed using the quasi-likelihood method in `edgeR`[4], where the dispersion is modelled from the GLM deviance and thereby stabilized with empirical Bayes shrinkage, to stabilize the estimates in the presence of limited replication.

**Adaptation of Spatial FDR to neighbourhoods** To control for multiple testing, we adapt the Spatial FDR method introduced by *Cydar* [3]. The Spatial FDR can be interpreted as the proportion of the union of neighbourhoods that is occupied by false-positive neighbourhoods. This accounts for the fact that some neighbourhoods are more densely connected than others. To control spatial FDR in the k-NN graph, we apply a weighted version of the Benjamini-Hochberg (BH) method. Briefly, to control for FDR at some threshold $\alpha$ we reject null hypothesis $i$ where the associated p-value is less than the threshold

$$\max_i p_{(i)} : p_{(i)} \leq \alpha \frac{\sum_{l=1}^{i} w_{(l)}}{\sum_{l=1}^{n} w_{(l)}}$$

Where the weight $w_{(i)}$ is the reciprocal of the neighbourhood connectivity $c_i$. As a measure of neighbourhood connectivity, we use the Euclidean distance to the kth nearest neighbour of the index cell for each neighbourhood.

#### 1.2 Accounting for batch effects

Comparing biological conditions often requires acquiring single-cell data from multiple samples, sometimes generated with different experimental conditions or protocols. This commonly introduces batch effects, which can have a substantial impact on the data composition and subsequently the topology of any k-NN graph computed across the single-cell data. Consequently, this will have an impact on the ability of *Milo* to resolve genuinely DA subpopulations between experimental conditions. To minimize the emergence of false positive and false negative results that might be introduced by such technical effects, we recommend the following procedure: (1) Batch effects should be mitigated with computational data integration. Defining the best tool for this task is beyond the scope of this work (a large number of integration methods have been reviewed and benchmarked in [5–7]). However, users should consider the type of output obtained by their integration method of choice, which can be a corrected feature space, a joint embedding or an integrated graph. Using a methods that produces a graph (e.g. BBKNN [8], Conos [9]) will lead to suboptimal results in DA testing with *Milo* as the refined neighbourhood search procedure will still be affected by the batch effect, as this relies on finding neighbors in PCA space. (2) Known technical or nuisance covariates should be introduced in the experimental design of DA testing, exploiting the flexible GLM framework used by *Milo*.

To demonstrate the use of including both steps in DA testing analysis, we simulated data for a continuous trajectory with two technical batches (Suppl.Fig.2A) and simulated a subpopulation of differential abundance between two conditions (Suppl.Fig.2B). We tested for DA between conditions on the k-NN graph generated before and after batch correction using the Mutual Nearest Neighbors (MNN) method [10]. In both the corrected and uncorrected data, we examined the effect of accounting for the batch in the experimental design of the GLM (`design = ~ batch + condition`). We find that, while alone MNN correction increases the true positive rate, accounting for the nuisance covariate in the testing design significantly increases accuracy and power both in the uncorrected and corrected graph (Suppl. Fig. 2C-D). The best DA prediction performance was achieved combining batch correction and an explicit experimental design adjusting for such nuisance covariates (Suppl. Fig. 2D).

To demonstrate how these results apply to a real dataset with a complex experimental design, we used a single-cell atlas of mouse gastrulation [11]. These data consist of 116312 cells collected from 411 embryos in 36 samples, across 10 developmental stages from E6.5 to E8.5 (Suppl. Fig. 3B). The samples were sequenced in 3 batches (Suppl. Fig. 3A). We used *Milo* to test for DA across developmental time points, on the k-NN graph built before and after batch correction with MNN. We find that including the sequencing batch in the DA testing design matrix removes false positives in the uncorrected graph, while increasing the number of detected DA neighbourhoods in the MNN corrected graph (Suppl. Fig.3C).

Taken together these results illustrate that while *Milo* can be used to robustly account for batch effects during DA testing, optimal results are achieved after the removal of major batch effects that are not readily accounted for by inclusion as a covariate in a linear model.

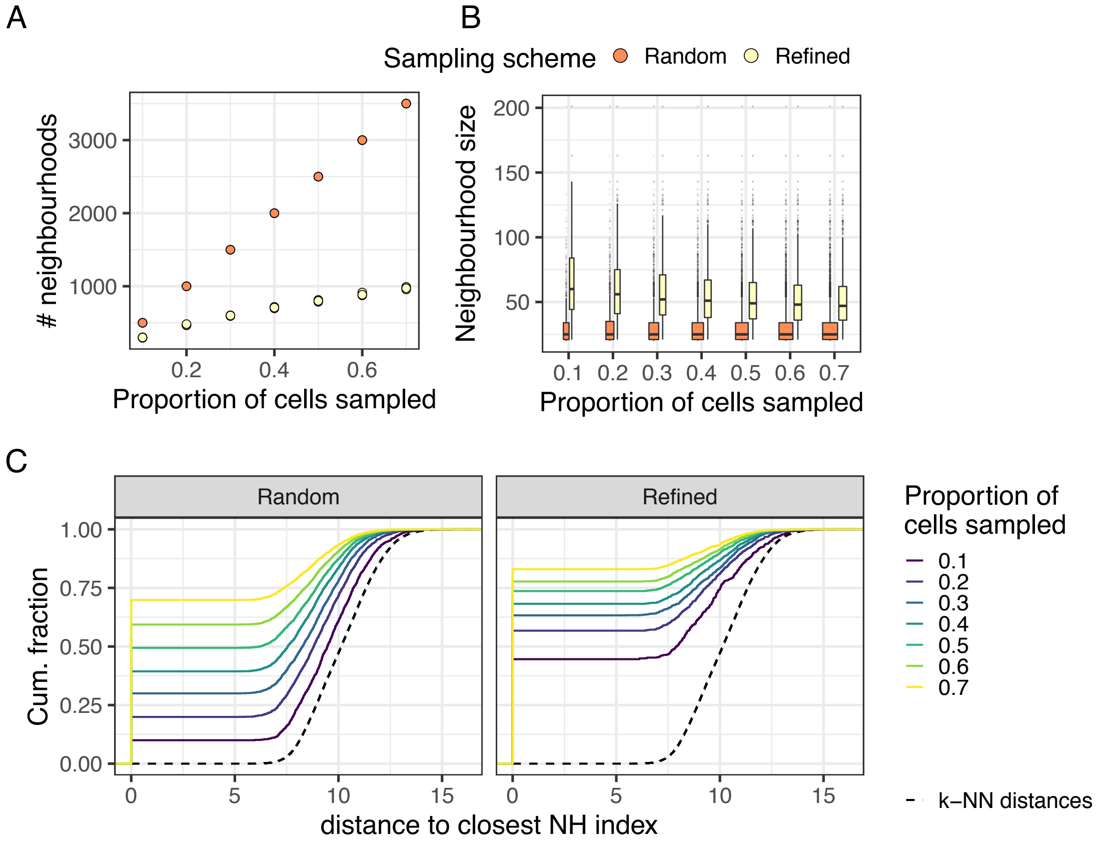

Supplementary Figure 1: **Random sampling of k-NN graph vertices is suboptimal compared to sampling with refinement.** (A) Sampling with refinement leads to selection of fewer neighbourhoods (B) Sampling with refinement leads to selection of bigger neighbourhoods for DA testing, independently of the initial proportion of cells sampled (C) Sampling with refinement generates robust neighbourhoods across initializations: for each index cell we calculate the distance from the closest index in a sampling with different initialization. The cumulative distribution of distances to the closest index is shown. The black dotted line denotes the distribution of distances between k nearest neighbors in the dataset (k=30) (NH: neighbourhood). Neighbourhood statistics were calculated using a simulated trajectory dataset of 5000 cells. All plots show results from three sampling initializations for each proportion.

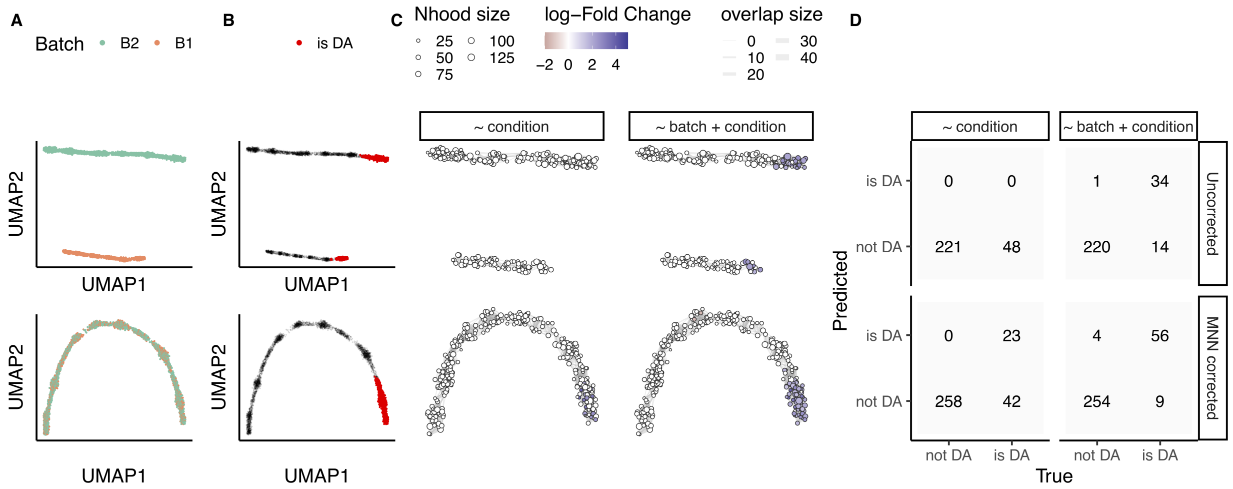

Supplementary Figure 2: **Tolerance of Milo to uncorrected batch effects in simulated data** (A-B) UMAP embeddings of simulated scRNA-seq data containing a batch effect, before batch correction (top row) and after correction with fastMNN (bottom row) (5000 cells). Cells are colored by simulated batch (A) and by presence of differential abundance between 2 simulated conditions (20% cells in condition 'A', 80% cells in condition 'B') (B). (C) A graph representation of the results from Milo differential abundance testing. Neighbourhoods were tested for DA between conditions, with (~ batch + condition) or without (~ condition) accounting for the simulated batch. Nodes are neighbourhoods, coloured by their log fold change between conditions. Non-DA neighbourhoods (FDR 10%). Node sizes correspond to the number of cells in a neighbourhood. Graph edges depict the number of cells shared between adjacent neighbourhoods. (D) Confusion matrices comparing the number of true and predicted DA neighbourhoods, with different batch effect correction (rows) and different testing design (columns).

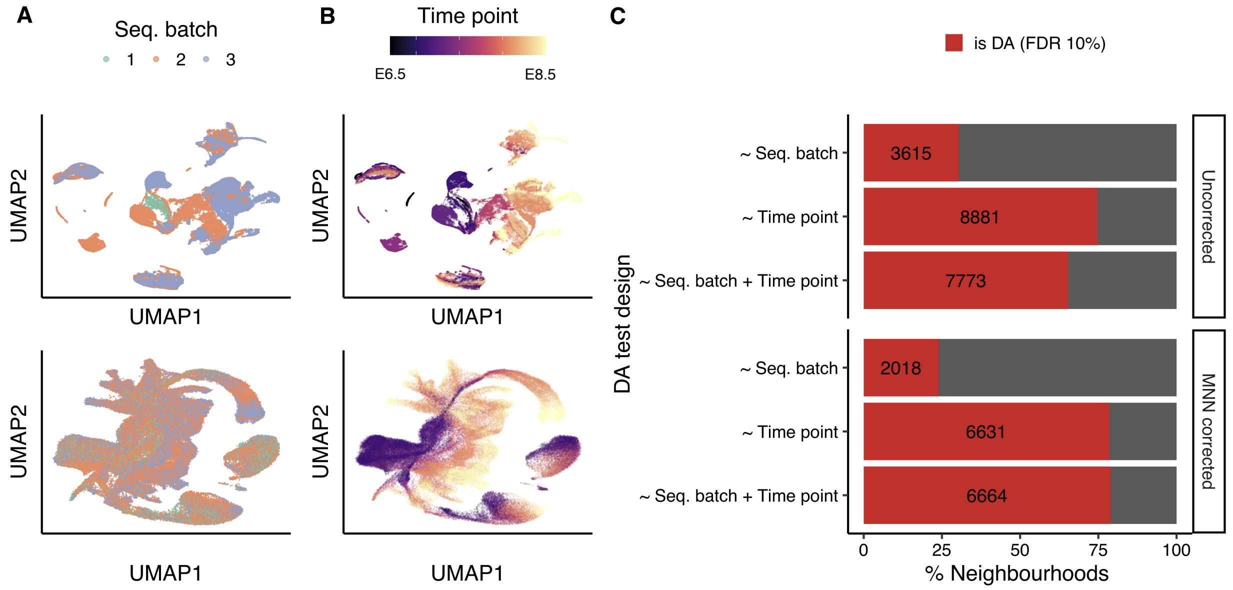

Supplementary Figure 3: **Tolerance of Milo to uncorrected batch effects in mouse gastrulation atlas** (A-B) UMAP embedding of mouse gastrulation atlas before batch correction (top row) and after correction with fastMNN (bottom row). Cells are colored by sequencing batch (A) and developmental time point (B). (C) Barplot depicting the percentage of DA neighbourhoods at FDR 10%, testing for different experimental covariates: DA between sequencing batches (∼ Seq. batch), DA across developmental time points (∼ Time points), DA across developmental time points accounting for the sequencing batch (∼ Seq. batch + time points). The total number of DA neighbourhoods is shown in each bar.

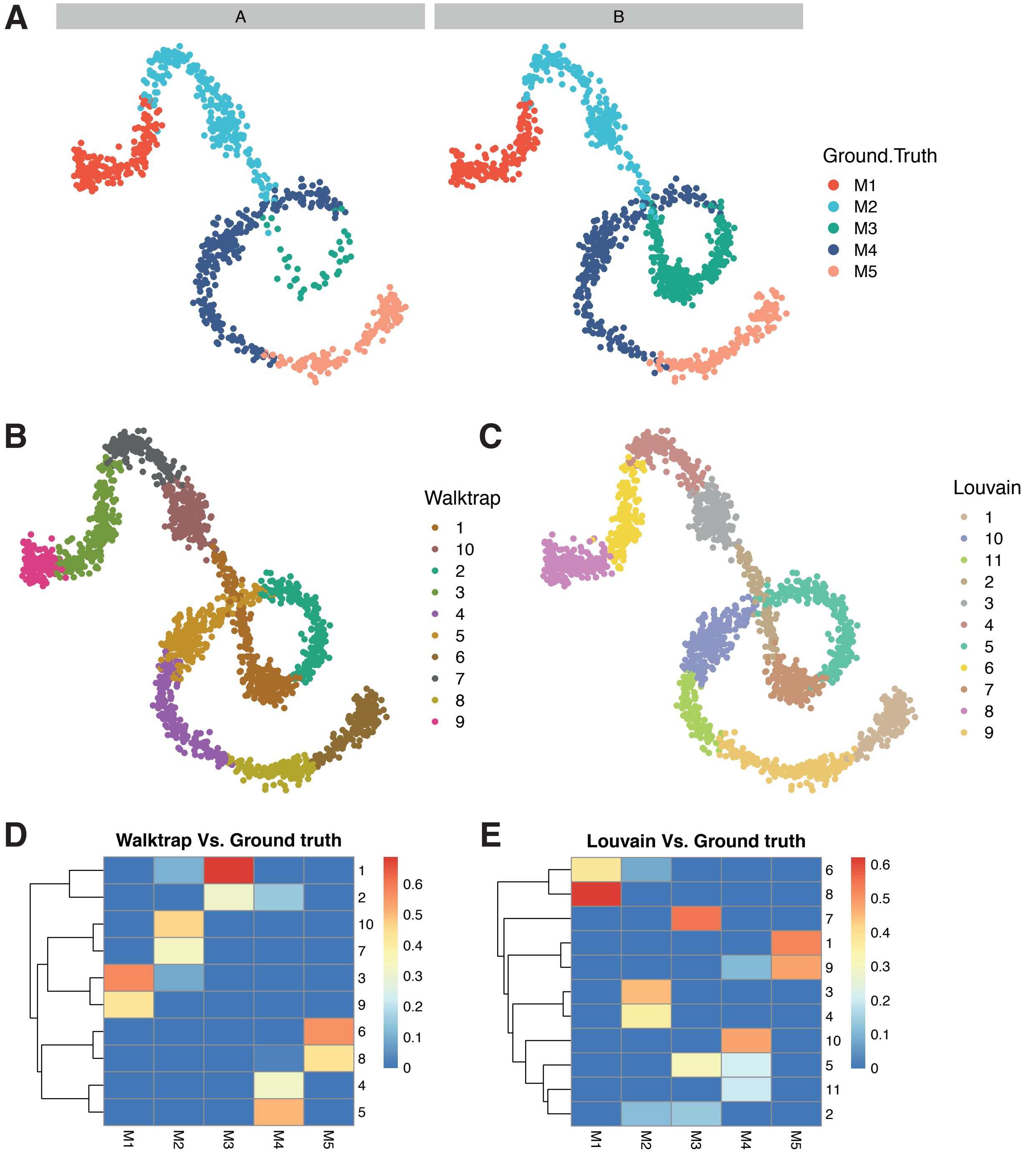

Supplementary Figure 4: **Graph-clustering does not faithfully capture simulated groups and differentially abundant subpopulations in a simulated continuous trajectory.** (A) A simulated linear trajectory of 2000 single-cells generated from 5 different groups, with cells assigned to either condition 'A' (left) or condition 'B' (right). (B) A Walktrap clustering of the data in (A) using the same k-NN graph. Cells are coloured by Walktrap cluster identity. (C) A Louvain clustering of the data in (A) using the same k-NN graph. Cells are coloured by the Louvain clustering identity. (D-E) Heatmaps comparing the numbers of cells in each cluster with respect to the ground truth groups in (A). Each cell is coloured by the proportion of cells from the column groups (ground truth) that are assigned to the respective cluster.

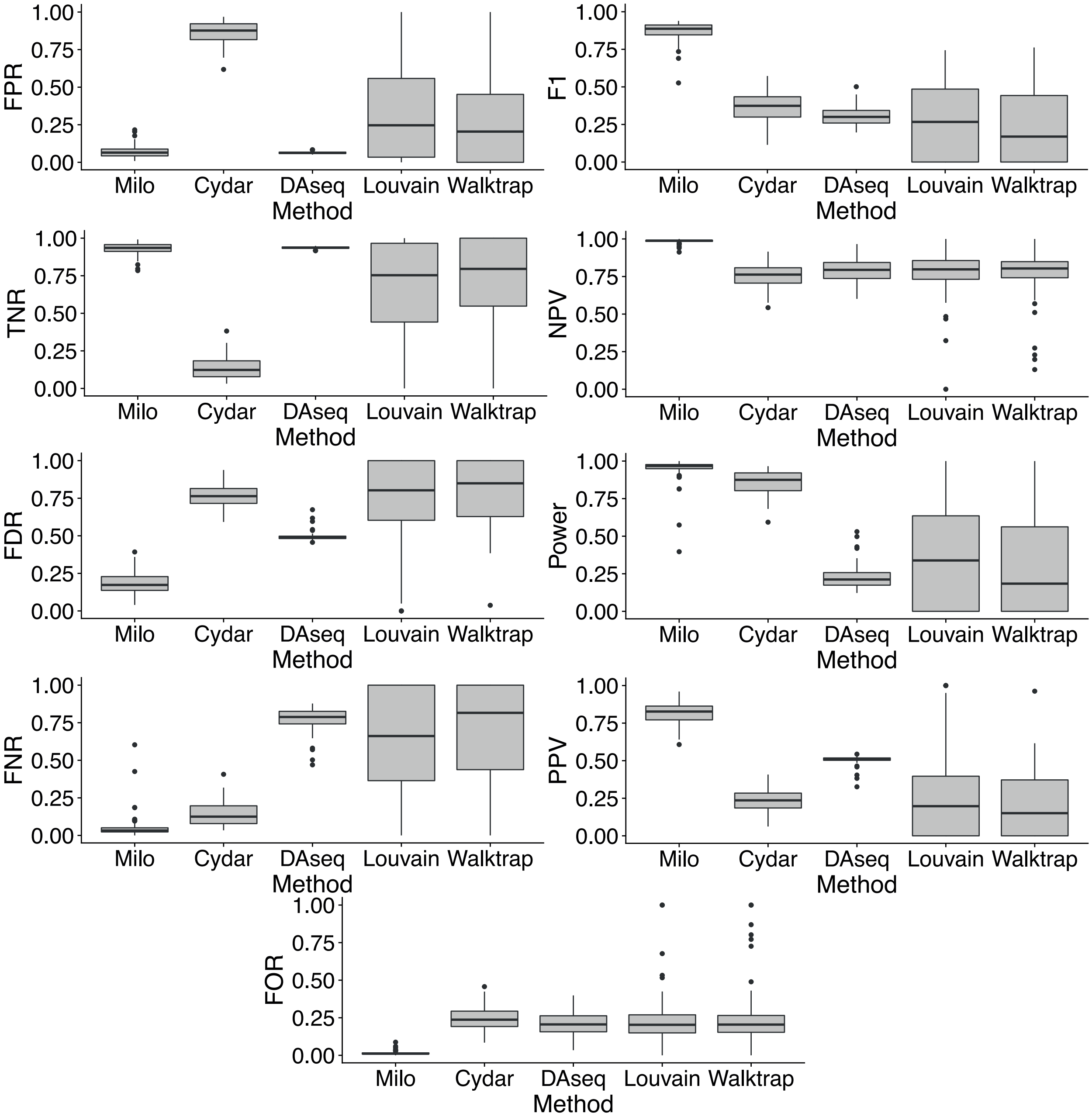

Supplementary Figure 5: **Comparison of Milo to alternative differential abundance methods** Each panel shows a measure of method performance computed across 100 independent simulations. Boxplots denote the median and interquartile range (IQR), with whiskers extending 1.5 x IQR; outliers are shown as individual points. Each analysis on each independent simulation used the same parameter values in Supplementary Table 1.

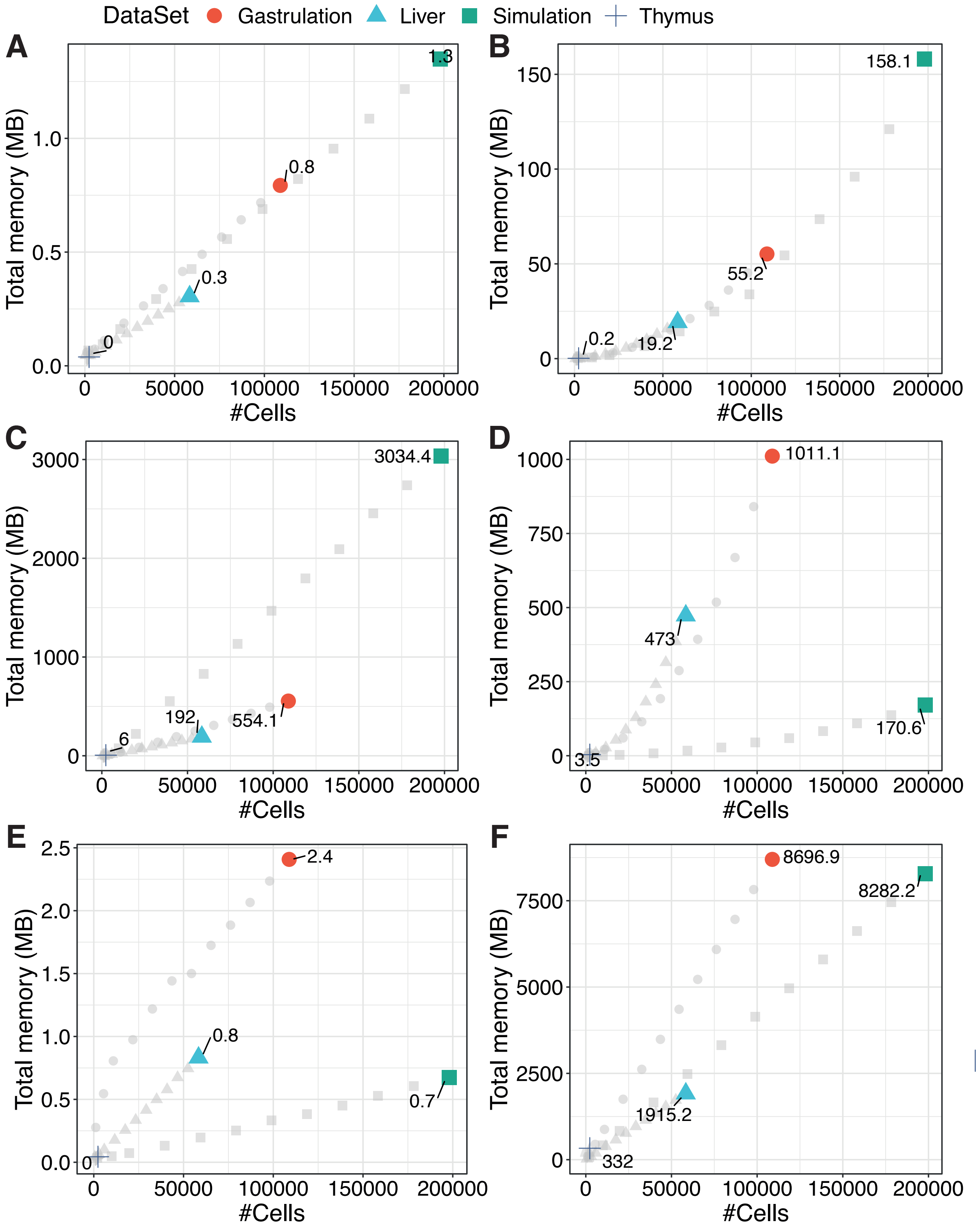

Supplementary Figure 6: **Memory usage across the Milo analysis workflow** Total memory usage across the steps of the Milo analysis workflow in 4 data sets containing different numbers of cells (Gastrulation: circles, Liver: triangles, Thymus: crosses, Simulation: squares). Grey points denoted down-sampled datasets of the corresponding type. Coloured points denote the total number of cells for the respective dataset. Total memory usage (y-axis) is shown in megabytes (MB). (A) k-NN graph building, (B) neighbourhood sampling and construction, (C) within-neighbourhood distance calculation, (D) cell counting in neighbourhoods according to the input experimental design, (E) differential abundance testing, (F) total in memory R object size. A fixed value was used in all data-sets for graph building and neighbourhood construction (k=30).

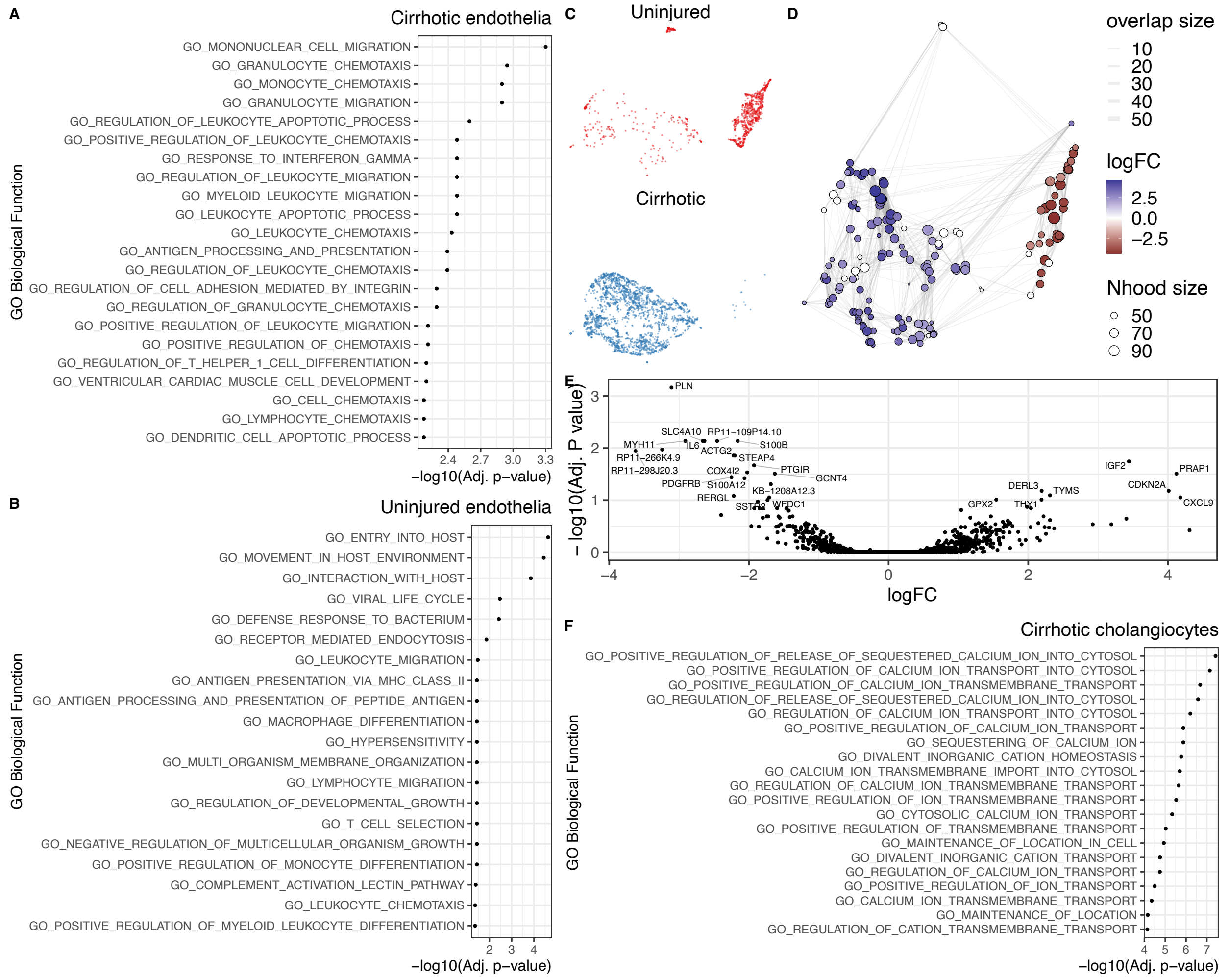

Supplementary Figure 7: **Downstream analysis of disease-specific subpopulations in liver cirrhosis** (A) GO term enrichment analysis on marker genes of cirrhosis-enriched endothelia. The top 20 significant terms are shown. (B) GO term enrichment analysis on marker genes of healthy-enriched endothelia. The top 20 significant terms are shown. (C-D) UMAP embedding (C) and graph representation (D) of neighbourhoods of 3369 cells from cholangiocytes lineage. (E) Volcano plot for DGE test on cholangiocytes DA subpopulations: the x-axis shows the log-fold change between expression in cirrhotic and healthy cholangiocytes. The y-axis shows the adjusted p-value. (F) GO term enrichment analysis on marker genes of cirrhosis-enriched cholangiocytes. The top 20 significant terms are shown.

### 1 Supplementary tables

| Method | Key Parameters | Values | Hypothesis testing |
|---|---|---|---|
| Milo | K | 10 | Negative binomial GLM, 10% FDR |
| | d | 15 | Negative binomial GLM, 10% FDR |
| Cydar | r | 2 | Negative binomial GLM, 10% FDR |
| DAseq | k.vector | 5-500, steps of 50 | Logistic classifier prediction, top 10% |
| Louvain + edgeR | K | 10 | Negative binomial GLM, 10% FDR |
| | d | 15 | |
| Walktrap + edgeR | K | 10 | Negative binomial GLM, 10% FDR |
| | d | 15 | |

Supplementary Table 1: **Method comparison parameter values.**
